# Left planum temporale growth predicts language development in newborns with congenital heart disease

**DOI:** 10.1101/439711

**Authors:** Andras Jakab, Eliane Meuwly, Maria Feldmann, Michael von Rhein, Raimund Kottke, Ruth O’Gorman Tuura, Beatrice Latal, Walter Knirsch, on behalf of the Research Group Heart and Brain

**Author notes:** **BGR**: brain growth rate, **CCS**: Bayley-III Cognitive Composite Score, **CHD**: congenital heart disease, **DBM**: deformation based morphome-try, **d-TGA**: dextro-transposition of the great arteries, **MCA:**middle cerebral artery, **MCS**: Bayley-III Motor Composite Score, **LCS**: Bayley-III Language Composite Score, **PMCC**: product-moment correlation co-efficient, **PCA:**posterior cerebral artery, **PT**: planum temporale, **TFCE**: threshold-free cluster enhancement, **SES**: socioeconomic status.

## Abstract

Congenital heart diseases (CHD) are the most common congenital anomalies, accounting for a third of all congenital anomaly cases. While surgical correction dramatically improved survival rates, the lag behind normal neurodevelopment appears to persist. Deficits of higher cognitive functions are particularly common, including developmental delay in communication and oral-motor apraxia. It remains unclear whether the varying degree of cognitive developmental delay is reflected in variability in brain growth patterns. To answer this question, we aimed to investigate whether the rate of regional brain growth is correlated with later life neurodevelopment. 44 newborns were included in our study, out of whom 33 were diagnosed with dextro-transposition of the great arteries (d-TGA) and 11 with other forms of severe CHD. During the first month of life, neonates underwent corrective or palliative cardiovascular bypass surgery, pre- and postoperative cerebral MRI were performed 18.7 ± 7.03 days apart. MRI was performed in natural sleep on a 3.0T scanner using an 8-channel head coil, fast spin-echo T2-weighted anatomical sequences were acquired in three planes. Based on the principles of deformation based morphometry, we calculated brain growth rate maps that reflected the rate of non-linear deformation that occurs between pre- and post-operative brain images. An explorative, whole-brain, threshold-free cluster enhancement analysis revealed strong correlation between the growth rate of the left planum temporale and the posterior operculum of the left frontal lobe and language score at 12 months of age, corrected for demographic variables (p=0.018, t=5.656). No significant correlation was found between brain growth rates and motor or cognitive scores. Post hoc analysis showed that the length of hospitalization interacts with this correlation with longer hospitalization stay results in faster enlargement of the internal cerebro-spinal fluid spaces. Our study provides evidence to the early importance of left-dominant perisylvian regions in language development even before the direct postnatatal exposure to native language. In CHD patients, the perioperative period results in a critical variability of brain growth rate in this region, which is a reliable neural correlate of language development at one year of age.

## Background

Late gestational and early postnatal neurodevelopment comprises important developmental processes, such as the rapid increase of cerebral volume and maturation of white matter. There is compelling evidence that congenital heart defects (CHD) adversely affect neurodevelopment during this period. However, the etiology of such cerebral injury is most likely multifactorial and not fully understood ^[1]^. As consequence of altered cerebral development, cognitive development may be impaired in children with more severe forms of CHD ^[2–5]^. Furthermore, CHD can lead to a variety of neurodevelopmental impairments: in infancy, they are characterized by abnormalities of muscular tone, feeding difficulties and developmental delays in major motor milestones ^[6]^, a variety of neurodevelopmental deficits manifest at later ages and may persist into adolescence and adulthood ^[7]^. Higher cognitive functions are particularly affected during later development ^[8, 9]^, and a developmental delay in communication and oral-motor apraxia also occur with a high prevalence ^[10–12]^. CHD children often score in the low-normal range on language tests, with 25% classified as at risk for both expressive and receptive language difficulties ^[13]^. Chronic hypoxemia due to cyanotic cardiac defects is more strongly associated with dysfunction of speech and language ^[14]^. The early identification of infants at risk for impaired neurocognitive development represents an important priority in clinical counselling of CHD patients leading to early therapeutic interventions.

Magnetic resonance imaging (MRI) studies have the potential to unravel early structural correlates of adverse neurocognitive outcome in CHD, such as smaller cerebral volumes *in utero*, and also during later life. Previous work found a correlation between cerebral volume prior to neonatal surgery and the degree of neurobehavioral development ^[3, 15]^. The global reduction of cerebral volume has been correlated with functional outcome ^[16]^, as the majority of studies could not unambiguously pinpoint a predilection of pathological changes, such as white matter injury ^[17–20]^, or delayed cortical gray matter development ^[21]^ to any of the cerebral lobes or smaller sub-divisions.

The additive effect of white matter injury and delayed cerebral maturation may explain why CHD infants lag behind normal neurodevelopment, and the varying degree of such injuries could presumably lead to variability in brain growth patterns among this population. Whether this variability in structural maturation is related to the different neurocognitive outcomes requires further evidence. Our purpose was to reveal possible morphological correlates of neurocognitive development, based on longitudinally acquired MRI in newborns who underwent corrective surgery for complex CHD. Most studies utilizing neuroimaging methods in CHD focused on a single time-point during pregnancy or child development ^[4, 16, 22]^, while others used repeated ultrasonography ^[23]^, MRI ^[20, 24]^or biometry such as head circumference measurement ^[18]^to characterize the impaired developmental trajectories. We assume that a single time-point approach may not be sensitive enough to characterize a critical period during which cerebral developmental is presumed to arrest or slowed. In contrast, repeated, longitudinal studies have the potential to link individual variability in postoperative cerebral growth rates to later cognitive development.

## Methods

### Patient population

Patients for the longitudinal MRI analysis were sampled from an ongoing prospective cohort study that investigates neurodevelopmental outcome in infants, who were operated for CHD within the first month after birth. Between December 2009 and August 2016, 78 infants with a severe type of CHD met the original study criteria. Inclusion criteria for the current work were (1) availability of pre- and post-operative MRI acquisitions with sufficient image quality, (2) availability of neurodevelopmental outcome assessment at one year of age.

Neonates with a suspected or confirmed genetic disorder or syndrome were excluded. 44 infants met these criteria and were included in the study.

Newborns were diagnosed with dextro-transposition of the great arteries (d-TGA, n=33) planned for biventricular repair, and non-TGA pathologies consisting of single ventricle undergoing staged palliation until Fontan procedure (n=5), interrupted aortic arch or coarctation of the aorta/transverse aortic arch hypoplasia (n=4), a case of complete unbalanced atrio-ventricular septum defect, one patient with perimembranous ventricular septum defect with open foramen ovale and one infant with pulmonary atresia with ventricular septum defect. Subjects were enrolled in the study after birth. Subject demographics are detailed in Table 1. Statistics on the presence of stroke, white matter injury or T2-hyperintensities in the pre- or postoperative images are summarized in Table S1.

**Table 1.** Demographic characteristics of the study population.

|  | d-TGA group | Other CHD | Total population | d-TGA – other CHD group difference |
| --- | --- | --- | --- | --- |
| Number of cases | 33 | 11 | 44 | - |
| Gender (M/F) | 24/9 | 10/1 | 34/10 | - |
| Gestational weight at birth, grams (mean ± SD, range) | 3436 ± 464 (2560 - 4270) | 3149 ± 312 (2775 - 3670) | 3364 ± 446 (2560 - 4270) | $p=0.029^*$ |
| Gestational age at birth, weeks (mean ± SD, range) | 39.75 ± 1.01 (37.3 - 41.4) | 39.41 ± 1.68 (37 - 41.7) | 39.66 ± 1.2 (37 - 41.7) | $p=0.54$ |
| Gestational age at first MRI, weeks (mean ± SD, range) | 40.53 ± 1.08 (37.6 - 42.1) | 41.52 ± 3.99 (38 - 51.6) | 40.78 ± 2.18 (37.6 - 51.6) | $p=0.43$ |
| Time between 1<sup>st</sup> and 2<sup>nd</sup> MRI scan, days (mean ± SD, range) | 18.7 ± 7.03 (7 - 38) | 17.82 ± 6.37 (12 - 33) | 18.48 ± 6.81 (7 - 38) | $p=0.70$ |
| Socioeconomic status | 9.42 ± 2.17 (5 - 12) | 8.09 ± 1.58 (6 - 11) | 9.09 ± 2.1 (5 - 12) | $p=0.038^*$ |
| Days of hospitalization | 31.12 ± 8.64 (13 - 47) | 37.36 ± 17.04 (19 - 73) | 32.68 ± 11.43 (13 - 73) | $p=0.27$ |
| Bayley CCS | 109.7 ± 12.37 (85 - 130) | 101.82 ± 18.48 (60 - 130) | 107.73 ± 14.32 (60 - 130) | $p=0.21$ |
| Bayley LCS | 92.72 ± 14.34 (65 - 132) | 95.73 ± 13.61 (74 - 118) | 93.49 ± 14.076 (65 - 132) | $p=0.54$ |
| Bayley MCS | 96.48 ± 11.75 (76 - 130) | 85.64 ± 15.71 (46 - 107) | 93.77 ± 13.51 (46 - 130) | $p=0.05$ |

\* significant at level $p<0.05$

The newborns underwent the following types of corrective surgeries: Arterial switch or Rastelli operation for patients with dextro-transposition of the great arteries, Norwood-type stage I palliation for patients with hypoplastic left heart syndrome and other forms of univentricular disease with hypoplastic aortic arch, and complex aortic arch reconstruction for patients with interrupted aortic arch or coarctation of the aorta/transverse aortic arch hypoplasia as well as corrective cardiac surgery.

The parents gave informed consent and the study has been approved by the local ethical committee, and research was conducted according to the principles expressed in the Declaration of Helsinki.

### MRI acquisition protocols

Neonatal cerebral MRI was performed in natural sleep on a 3.0T clinical MRI scanner using an 8-channel head coil, and fast spin-echo T2-weighted anatomical sequences were acquired in axial, sagittal and coronal planes. The sequence parameters for the anatomical MRI were the following. TE/TR: 97/5900, flip angle: 90, pixel spacing: 0.35 * 0.35 mm, slice thickness: 2.7 mm. During the recruitment, the MRI scanner’s software was upgraded; 25 cases were scanned before upgrade, but pre-and post-operative scans were performed on the same scanner software level for all patients. Newborns underwent the first, preoperative MRI at the corrected gestational age of 40.77 ± 2.18 (37.6 – 51.6) weeks, and the postoperative follow-up was performed at 43.52 ± 2.33 (40 – 54) weeks; the time difference between the two MRI exams were 18.48 ± 6.81 (range: 7 – 38) days.

### Image processing

Fig. 1 gives an overview of the image analysis steps used in the study. MR images were first masked using a semi-automated approach to remove voxels corresponding to non-brain tissue in the Slicer 3D software. Then, the axial, sagittal and coronal T2-weighted images were reconstructed to a 3D image with isotropic voxel spacing of 1 mm using the BTK toolkit (Rousseau *et al.*, 2013).

To capture longitudinal changes, we employed a modified version of deformation-based morphometry, in which the anatomical deformation that matches the pre- and post-operative images are calculated on the case level, and the resulting deformation field is standardized across the population.

**Figure 1.**
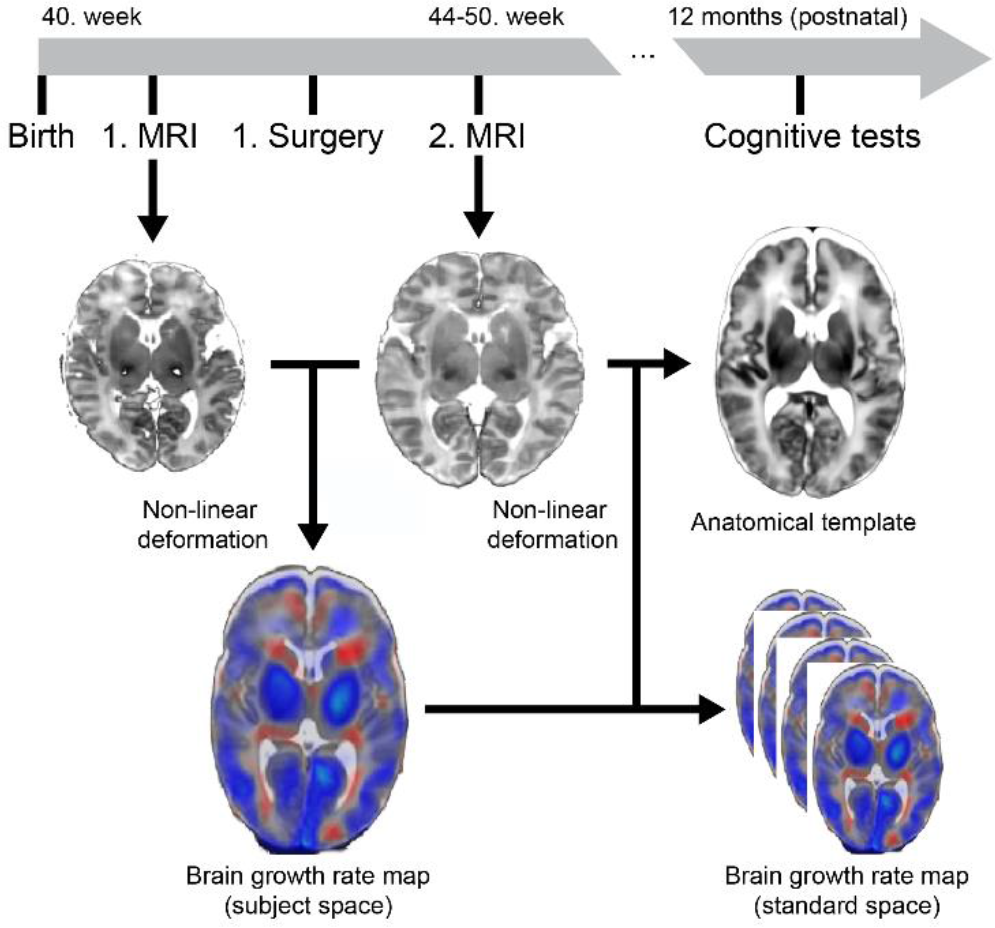
Overview of the image acquisition and analysis steps.

This was carried out using the following steps: (1) rigid alignment with 6 degrees of freedom of the pre-operative, brain-extracted T2-weighted image to the post-operative MRI, (2) calculating the deformation field that matches the preoperative image with the postoperative MRI based on a high-resolution freeform non-linear deformation algorithm (implemented in the NIFTIREG tool (Modat *et al.*, 2010), reg_f3d command, control grid size: 5 mm isotropic, weight of the bending energy penalty term: 0.005, gradient smoothing with a kernel of 4 mm), (3) transforming the natural logarithm of the Jacobian determinant of this transformation to brain growth rate (BGR) using the following equation:

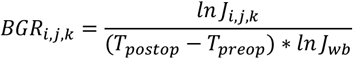

*where J_i,j,k_ is the Jacobian determinant of the pre- to postoperative deformation field at voxel i,j,k, J_wb_ is the average Jacobian determinant over the brain tissue*, (*T_postop_* - *T_preop_*) *is the time difference between the post- and preoperatively acquired MR images.*

We then (4) co-registered the preoperative neonatal brains to a standard, 40. gestational (term) week image template, (5) transformed the BGR maps using the registration step (4) to the 40. gestational week template, and (6) performed spatial smoothing with a Gaussian kernel of σ=5 mm.

### Neurodevelopmental testing

Neurodevelopmental testing was performed using the Bayley Scales of Infant and Toddler Development, Third Edition (Bayley-III), at 12 months of age with three developmental domains: cognitive (CCS), language (LCS) and motor composite standard scores (MCS) with a mean score of 107.73, 93.5 and 93.77 in our cohort, respectively.

### Statistical tests

Descriptive statistics was performed using an independent two-sample t-test assuming equal variance. Statistical analyses on the whole-brain BGR maps were performed using the FSL software package (Jenkinson *et al.*, 2012). Linear regression analysis was performed to evaluate the association of BGR with the 12 month Bayley-III scores. Multiple comparison correction of the regression analyses was performed using threshold-free cluster enhancement (TFCE) (Smith and Nichols, 2009), with randomized nonparametric permutation testing implemented in the *randomise* program (Winkler *et al.*, 2014), version 2.9, part of FSL (build 509). The default parameter settings of TFCE with 5000 permutations were used, and results were restricted to a brain mask in template space corresponding to the brain parenchyma. For the initial analysis, family-wise error corrected p≤0.025 was accepted as significant, this threshold accounted for the fact that positive and negative correlations were investigated separately. We chose to correct the first model for a variable that is known to correlate with MRI appearance and hence may influence DBM results: “MRI software version” (Shuter *et al.*, 2008), since the scanner was upgraded during the recruitment period of the subjects.

Furthermore, two known variables that may have influenced the neurocognitive tests were also included: parental socioeconomical status (SES) and gender (Largo *et al.*, 1989; Wickremasinghe *et al.*, 2012; Ronfani *et al.*, 2015). The dichotomous variable “MRI software version” was used to define two exchangeability blocks, and observations were permuted only within blocks. To demonstrate the effect size of linear regression analysis, maps depicting the Pearson product-moment correlation coefficient in Matlab R2014 (Mathworks Inc., Mattick, USA) were calculated. Next, post hoc analysis was performed using stepwise linear regression in IBM SPSS V22 (IBM, Armonk, New York) to select any further demographic or clinical parameters that could influence the correlation between regional BGR and the Bayley-III scores. During the stepwise selection of variables from a pool of 21 demographic and clinical parameters (Supplementary Table 2), the probability of F was set to 0.05 for a variable to enter and 0.1 for removal. Finally, a second linear regression analysis with randomized nonparametric permutation testing was performed over the whole brain, which was corrected for the confounding variables found in the post hoc analysis step, and TFCE-corrected p≤0.05 was accepted significant.

## Results

### Correlation between regional brain growth rate and 12 month Bayley-III composite scores

We observed positive BGR values over the entire brain parenchyma, reflecting increasing brain volumes during the perioperative period. Relative shrinkage was seen at the interface of the brain parenchyma and cerebro-spinal fluid interfaces supratentorially, which was most likely the consequence of the mildly increasing ventricular dilatation.

**Figure 2.**
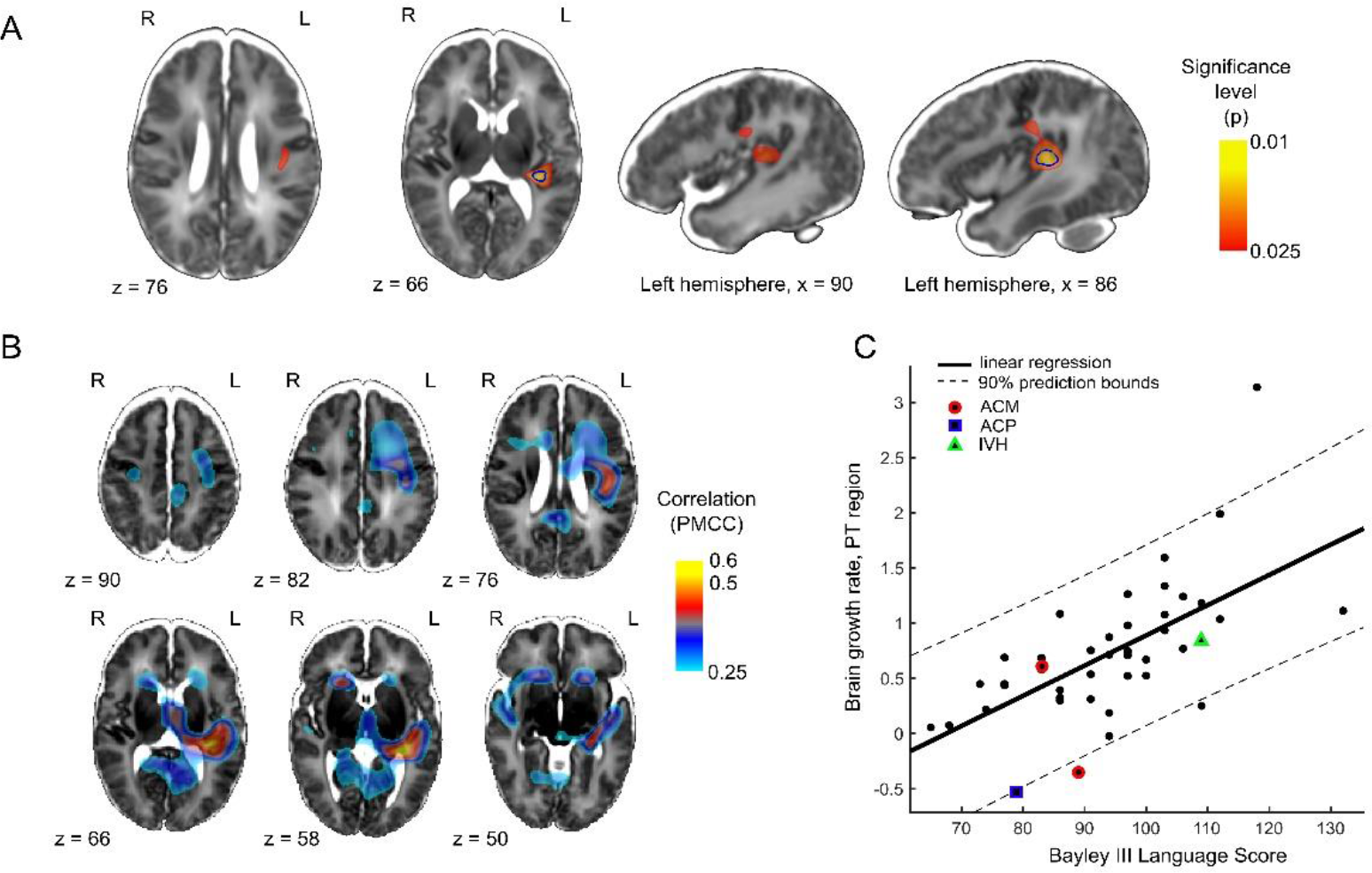
Correlation of perioperative regional brain growth rate with the one-year Bayley-III Composite Language Score at 12 months of age. (A) brain regions significantly (TFCE-corrected p≤0.025) correlating with LCS are shown as red-to yellow clusters overlaid on a T2-weighted MRI template, blue outline (within the red overlay): significant cluster after adjusting for length of hospital stay, (B) the correlation coefficient map (thresholded at R≥0.25), (C) correlation plot. PMCC: Pearson’s product-moment correlation coefficient. ACM: two patients with middle cerebral artery stroke, ACP: one patient with posterior cerebral artery stroke, IVH: one patient with intra-ventricular hemorrhage reported on the pre-operative MRI.

Whole-brain explorative analysis revealed that BGR of the left posterior perisylvian region was significantly positively correlated with the LCS at 12 months of age; no other clusters of correlation survived the statistical significance threshold. This effect was localized to a circumscribed region comprising 1597 voxels (1.013 cm3) in one cluster, in which the peak TFCE-corrected significance (p=0.0094, maximum t=5.41, R=0.557) was localized to the left planum temporale. The cluster extended laterally from the border of the left lateral ventricle towards the outer third of the Heschl gyrus, respecting the gray matter / cerebrospinal fluid interfaces along the planum temporale (Fig. 2A). Superiorly, it extended into the white matter of the posterior parts of the left frontal operculum. A cluster of strong correlation between BGR and LCS (R>0.5) was co-localized with the planum temporale on the corresponding correlation coefficient maps, however, additional clusters of moderate to strong, but not significant correlation (0.25<R<0.5) were also found around the frontal and occipital horns of the lateral ventricles, and in the left frontal white matter (Fig, 2B).

No significant positive correlation surviving the correction threshold 0.025 was found between the BGR and the CCS (maximum of p=0.0475, t=4.15) or MCS (maximum of p=0.433, t=3.59). No significant correlations were found between the BGR maps and the CCS (maximum of p=0.13, t=3.77), the MCS (maximum of p=0.64, t=3.49) or the LCS (maximum of p=0.28, t=3.61). By qualitative inspection, the correlation coefficient map of the CCS resembled greatly that of the LCS (Fig. 3), and the correlation between CCS and LCS was strong (R=0.7249). CCS correlated with BGR more pronouncedly in midline regions in the thalamus, and bilaterally in the pre- and postcentral gyrus. The correlation between MCS and BGR showed more bilateral patterns and left frontal dominance, and weak correlation was found between LCS and MCS (R=0.1096). Contrary to our *a priori* hypothesis, BGR did not correlate with SES, gender or “MRI software version” when adjusting for the effect of LCS.

### Post hoc analysis of disease severity, clinical and demographic effects

We calculated the mean BGR in the cluster in the planum temporale (PT) acquired from the previous analysis step, and *post hoc* stepwise linear regression analysis was performed to automatically select the possible clinical and demographic variables that predict brain growth rate in the PT cluster in combination. Two variables emerged from the stepwise linear regression, the LCS and the days of hospital stay (Table 2).

**Table 2.** Post-hoc stepwise linear regression analysis of the correlations between the BGR in the planum temporale region, LCS and significant confounders.

| Variable | Included in *a priori* test? | Test statistic (t) | Significance (p) | Standardized coefficient (beta) | Unstandardized coefficient (B, CI) |
| --- | --- | --- | --- | --- | --- |
| LCS | Yes | 5.11 | $8 \times 10^{-7}$ | 0.575 | 0.025<br>(0.015 – 0.035) |
| Length of hospital stay | No | 2.89 | 0.0061 | 0.325 | 0.017<br>(0.005 – 0.029) |

Next, the effect of the length of hospital stay on the BGR over the whole brain was evaluated, controlling for the variability of LCS, SES, gender and MRI software version (Fig. 4). We found that length of hospital stay positively correlated with BGR independently from LCS in both hemispheres periventricularly, indicating excessive growth of CSF spaces and in the central parts of the white matter, the volume of the significant cluster was 8565 voxels (5.43 cm3), maximum of p=0.018, t=5.656 (Fig. 4A). Local volume reduction was associated with length of hospital stay over the entire brain surface; however, significance was not reached after TFCE correction (Fig. 4B). No further parameters related to disease severity contributed significantly to this model. Results of the linear regression for variables not selected are displayed in Supplementary Table 2.

Three subjects had ischemic lesions (two cases with arteria cerebri media, one case with arteria cerebri posterior stroke) and one intra-ventricular hemorrhage, the latter was detected on the preoperative MRI. Due to the small number of such cases, we decided not to include a variable for ischemic lesions as a confounder, but demonstrated the relationship between BGR and cognitive outcomes in these cases individually (Fig. 2C). The LCS and the planum temporale cluster’s growth rate was markedly lower in the three cases with ischemic lesions, two of them demonstrated a shrinkage of the planum temporale cluster in the postoperative MRI. The one case with intra-ventricular hemorrhage demonstrated average regional BGR in the planum temporale and a normal LCS score (109) at 12 months.

**Figure 3.**
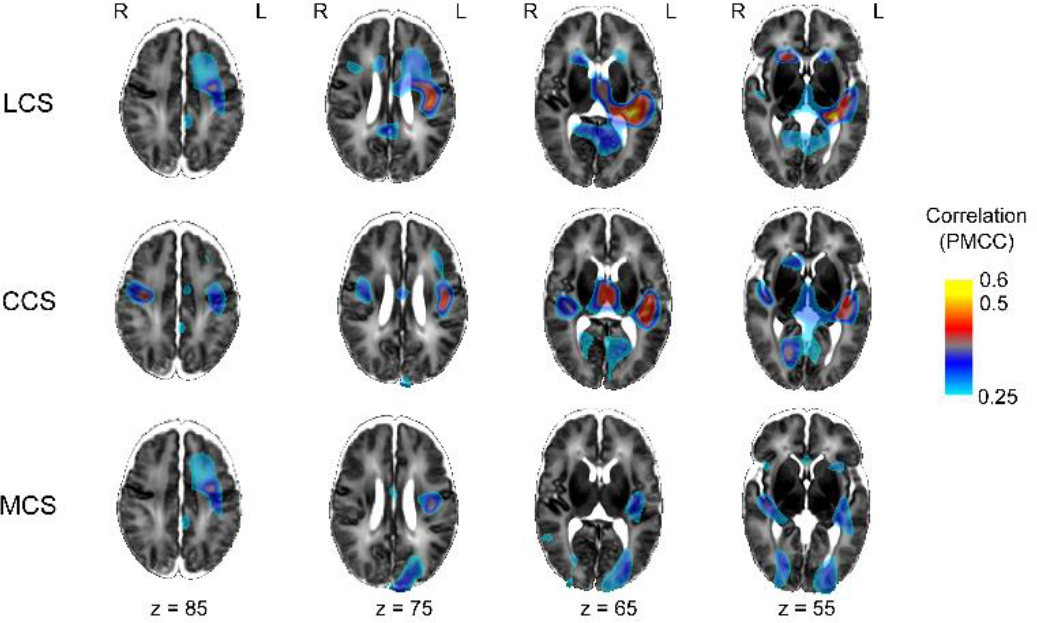
Comparison of brain growth rate correlating with the one-year Bayley-III cognitive, motor and language scores. Brain regions with moderate to strong (R≥0.25) positive correlation appear as a color-coded overlay fused with the MRI template. PMCC: Pearson’s product-moment correlation coefficient.

**Figure 4.**
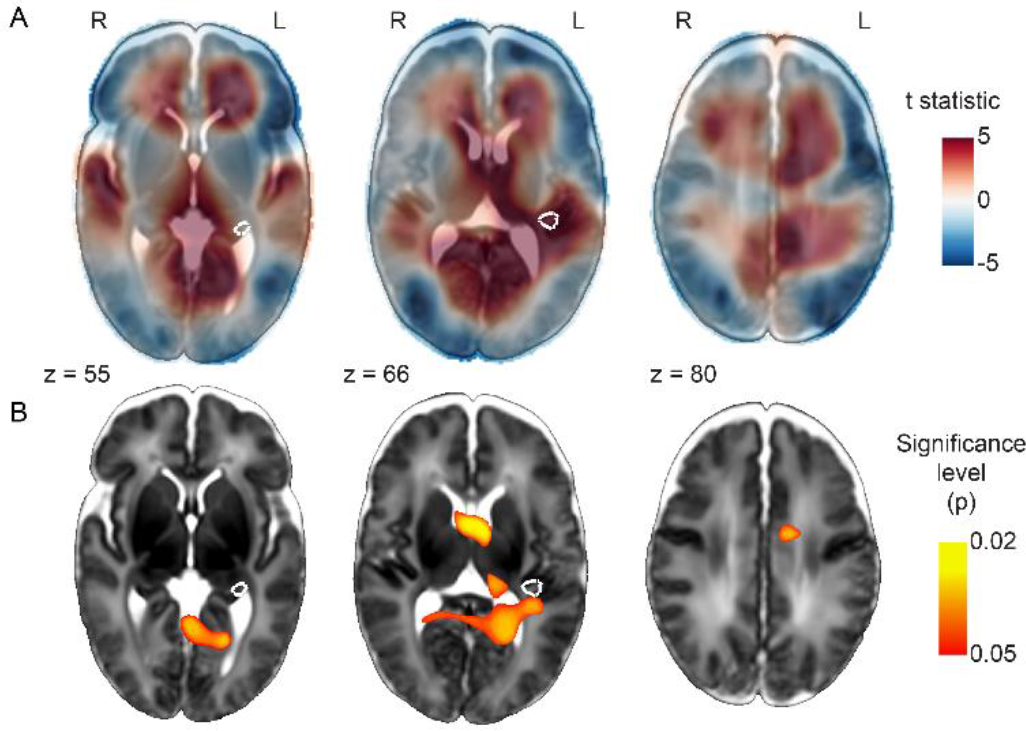
Effect of length of hospital stay on regional brain growth rate, corrected for LCS, SES, gender and MRI software version. (A) Brain regions with significant (TFCE-corrected p≤0.025) positive correlation are depicted as red to yellow overlays, and were fused with the MRI template. (B) Corresponding test statistic maps (t), positive values referring to positive correlation between regional growth rate and hospitalization duration. The brain region in which BGR correlated with LCS is displayed as white outlines.

### Correlation between regional brain growth and 12 month Bayley-III scores, corrected for length of hospital stay

A second linear regression analysis with randomized nonparametric permutation testing was performed, corrected for the length of hospital stay, SES, gender and MRI software version. BGR correlated positively with LCS after TFCE correction (p≤0.05) in the left hemispheric perisylvian region similar to the model not including length of hospital stay as a confounder. This well-circumscribed cluster comprised 861 voxels (0.546 cm3), peak TFCE-corrected significance was p=0.0268, maximum t=4.4. The modified model’s significant cluster is demonstrated in Fig. 2A as blue outlines.

## Discussion

CHD infants demonstrate global and regional cerebral growth within three weeks after cardiopulmonary bypass surgery and intensive care stay. While regional brain growth rates during this period showed no significant association with motor or general cognitive outcome, perioperative growth in the left-dominant perisylvian region was strongly correlated with language performance at one year of age. This close relationship was observed during the critical life period of CHD infants when they undergo the first cardiovascular bypass surgery for the underlying abnormality.

We also found evidence that the length of hospital stay contributes to the observed variability of brain growth and language outcome. In a multivariate model including gender, MRI software version and SES, regional brain growth rate of the left planum temporale and the posterior operculum of the left frontal lobe strongly correlated with language development at 12 months of age. Whereas this newly described morphological substrate for early language development appeared to be left lateralized and well-circumscribed, such anatomical specificity was not observed for the more general cognitive and motor development of the infants.

The regions in which regional growth moderately correlated with CCS comprised the bilateral pre- and postcentral gyri, the medial thalamus, posterior, the medial occipital white matter and the posterior horn of the lateral ventricle. This may reflect that CCS and MCS scores are more generalized markers of neurodevelopment, with more distributed neuroanatomical substrates and less pronounced morphology-function link at such an early age.

The length of hospital stay showed correlation with brain growth rate in areas around the CSF spaces, which could reflect increasingly dilated lateral ventricles or volume loss of the cerebral white matter surrounding the ventricles. Length of hospital stay is a variable correlating with postoperative disease severity. It can be influenced by low cardiac output syndrome including low cerebral perfusion, longer need for inotropic support, and longer time for intubation. Other factors such as infections, rhythm disorders, renal insufficiency and higher nutritive demands my affect cerebral growth. Dilatation of lateral ventricles have been shown in older children at 2-3 years of age before Fontan procedure correlated with adverse neurodevelopmental outcome ^[25]^.

Our results are in agreement with previous works that found a link between left perisylvian morphology and language development in high-risk infants, such as in newborns suffering from hypoxic-ischemic injury ^[26]^ or preterm infants ^[27]^. In CHD, white matter volume was found to correlate with language development, but not broader developmental indices ^[28]^, similarly to our measurements, and in concordance with studies on extremely preterm infants ^[29]^. While these studies establish a link between morphological features at an early stage of neurodevelopment and later life cognitive impairment, it is not yet clear if this is a consequence of an underlying pathology associated with CHD or reflects inter-individual variability in normal development. Our study sample consists of infants with sub-normal and normal cognitive scores and therefore represents a population with impaired development rather than the normal variability.

Our findings provide evidence that regional brain growth rate within the first weeks of life predicts language processing in infants. We have demonstrated that the brain growth rate in the left planum temporal and left posterior frontal operculum during the first three weeks of life is predictive of later language performance in CHD. This marks a high anatomical specificity of language processing immediately after birth, providing further evidence that language processing in infants is supported by neuroanatomical structures similar to the adult brain. The left superior temporal and angular gyri are already active in infants during speech processing at 3 months of age ^[30–33]^. These functional neuroimaging studies suggest that language processing is already localized to the left perisylvian region shortly after birth – or even before. The pre-existing hemispheric asymmetry of the perisylvian region most likely underlies the functional specialization for language at such an early age: it is known that neonates are born with left dominant planum temporale and deeper right superior temporal sulcus ^[34–36]^, and the development of language structures supporting processing areas also demonstrate hemispheric asymmetry in new-borns ^[37–40]^. These in utero and early postnatal findings imply strong genetic influence on the observed perisylvian asymmetry, however, it is also known that early life experience, such as exposure to language further shapes hemispheric lateralization, and contributes to the more pronounced lateralization patterns seen in later life ^[41, 42]^. Similarly, pathological processes can interfere with language development during infancy, and this leads to neuroplastic reorganization of language-processing areas in cases of structural damage to the left perisylvian region ^[43]^. Failure to develop left-hemispheric specialization may be the sign of maladaptive neurodevelopment, such as in autism spectrum disorders ^[44]^.

While surgical correction and palliation of CHD improved survival rates in CHD, the lag behind normal neurodevelopment appears to persist despite corrective surgery ^[45]^. The impairment of higher cognitive functions becomes most apparent during childhood or adolescence, hallmarking the importance of close neurodevelopmental follow-up. Single time-point studies confirmed a variety of neuroanatomical correlates of cognitive development in CHD, such as reduced global brain volumes ^[4, 16, 17, 20, 22, 24]^, cerebral white matter volume and microstructure ^[28, 46]^, and hippocampal volume reduction ^[47]^.

There is also a lack of evidence whether individual differences in early pre- or post-operative growth rates are useful predictors of cognitive outcome in complex CHD. Longitudinally acquired cerebral MRI offers the possibility to noninvasively characterize trajectories of structural brain development in children with complex CHD, and to find a link between growth and long-term cognitive development. Previous research found that cerebral growth rate following surgical correction may not be different in the first 3 months of life in a mixed group of complex CHD compared to typical development ^[20]^. On the contrary, it was shown that growth trajectories might exhibit differences depending on the underlying physiology ^[48]^. Correlation between growth rates and neurodevelopment was confirmed for other risk groups and pathologies. In preterms, somatic growth velocity during hospitalization was proven to be a possible independent predictor of adverse neurodevelopment ^[49]^. Impaired brain growth trajectories were associated with fetal alcohol spectrum disorders, which condition is also characterized by cognitive impairment ^[50, 51]^. Neurocognitive outcomes in high risk infants may, however, be better predicted by the multi-parametric analysis of brain morphology, brain connectivity and function ^[52, 53]^.

The following limitations of our study merit mentioning. We observed a remarkably large effect of brain growth over a rather short period of time – on average 3 weeks – on language outcomes. It is known that the maturation of white matter structure during the first months of life is particularly rapid in the cortico-thalamic, corticospinal fibers, as well as in subcortical areas ^[54]^, and that the morphological maturation of brain convolutions continue after birth. Based on this knowledge, we presume that variability of the left planum temporale growth contributed significantly to the differences seen in the language development, however, other factors, such as perioperative brain injury, the surgical technique and complications, such as ischemic lesions or intra-ventricular hemorrhage may influence the observed relationship between growth and outcome. We found no relationship between the presences of T2-hyperintensity, white matter damage and left planum temporale growth, however, we were not able to discern such link for ischemic brain lesions due to the low number of such cases (3/44).

The heterogeneity in our sample calls for careful attention when generalizing the results to all types of CHD, and long-term follow-up results are necessary to validate if the revealed morphology-outcome link is sustained into later stages of development. It remains an open question if a similar correlation between early neonatal brain growth in the left planum temporale and 1-year language outcome persists in other risk groups, such as in pre-terms or term born, normally developing infants. While the lack of controls hinders the more general interpretation of our results, we hypothesize that CHD would be an optimal model to study such correlations due to the higher variability of cognitive outcomes and growth rates in this population. The interpretation of our results is further limited by the fact that the utilized DBM method cannot distinguish between white matter and cortical growth, as the images were not segmented into such compartments prior to analysis. Despite this, DBM might be appropriate for the analysis of the newborn brain, as brain maturation results in rapid changes in the MRI appearance, rendering automatic tissue type classification difficult ^[55]^. In our study, we did not determine brain weight (corrected for body weight) as a result parameter in our study due to the specific analysis of very small regional parts of the brain.

Our study provides further evidence to the early importance of left-dominant perisylvian regions in language development even before the lasting exposure to native language. Inter-individual variability of regional brain growth rates in this area was found to be linked to variability in language performance at 12 months of age. Deformation based morphometry, based on longitudinal MRI before and after the corrective surgery for CHD may help to identify at risk-infants for impaired language development, and may facilitate better clinical counseling for high risk infants.

## Acknowledgements

A.J. was supported by the OPO Foundation, the FZK Foundation, the Stiftung für wissenschaftliche Forschung an der UZH and the EMDO Foundation, R.T. was supported by the EMDO Foundation, B.L. was supported by the Mäxi Foundation.

---

**Supplementary Table 1.** Summary of abnormalities reported on the pre- and postoperative MRI.

| Type of finding | Preoperative / postoperative |
| --- | --- |
| Intraventricular hemorrhage | 1/1 |
| Subdural hemorrhage | 15/16 |
| Plexus choroideus hemorrhage | 12/10 |
| Stroke, anterior cerebral artery | 0/0 |
| Stroke, middle cerebral artery | 2/2 |
| Stroke, posterior cerebral artery | 1/1 |
| T2 hyperintensity | 30/26 |
| White matter injury | 9/8 |

**Supplementary Table 2.** Post-hoc stepwise linear regression analysis of the correlations between the BGR in the planum temporale region, LCS, significant and non-significant confounders.

| Variable | Included in *a priori* test? | Included in *post hoc* test? | Test statistic (t) | Significance (p) | Unstandardized coefficient |
| --- | --- | --- | --- | --- | --- |
| LCS | Yes | Yes | 5.11 | 8*10<sup>-7</sup>* | 0.025 |
| Length of hospital stay | No | Yes | 2.89 | 0.0061* | 0.017 |
| Gestational age at birth | No | No | 1.942 | 0.059 | 0.209 |
| Gender | Yes | Yes | 0.719 | 0.477 | 0.081 |
| Socioeconomic status | Yes | Yes | -1.386 | 0.173 | -0.161 |
| Birth weight | No | No | 0.676 | 0.503 | 0.077 |
| Head circumference at birth | No | No | 0.166 | 0.869 | 0.019 |
| Apgar Score at 5 minutes | No | No | 1.172 | 0.248 | 0.130 |
| Anterior cerebral perfusion duration | No | No | 0.568 | 0.573 | 0.065 |
| Extra-corporal circulation duration | No | No | 0.605 | 0.549 | 0.071 |
| Age at first MRI | No | No | 1.292 | 0.204 | 0.143 |
| Age at second MRI | No | No | 0.903 | 0.372 | 0.103 |
| Univentricular pathology | No | No | -1.021 | 0.314 | -0.115 |
| Plexus choroideus bleeding. preoperative MRI | No | No | 0.013 | 0.989 | 0.002 |
| White matter injury. preoperative MRI | No | No | -0.044 | 0.965 | -0.005 |
| T2 hyperintensities. preoperative MRI | No | No | -1.394 | 0.171 | -0.154 |
| Subdural hemorrhage. preoperative MRI | No | No | 1.441 | 0.158 | 0.176 |
| Plexus choroideus bleeding. postoperative MRI | No | No | -1.185 | 0.243 | -0.137 |
| White matter injury. postoperative MRI | No | No | -1.184 | 0.244 | -0.137 |
| T2 hyperintensities. postoperative MRI | No | No | -0.761 | 0.451 | -0.087 |
| Subdural hemorrhage. postoperative MRI | No | No | -1.182 | 0.244 | -0.137 |

\* significant at level $p < 0.05$

